# Efficient and scalable integration of single-cell data using domain-adversarial and variational approximation

**DOI:** 10.1101/2021.04.06.438733

**Authors:** Jialu Hu, Yuanke Zhong, Xuequn Shang

**Affiliations:** School of Computer Science, Northwestern Polytechnical University, 127 West Youyi Rd., 710072, Shaanxi, China

## Abstract

Single-cell data provides us new ways of discovering biological truth at the level of individual cells, such as identification of cellular sub-populations and cell development. With the development of single-cell sequencing technologies, a key analytical challenge is to integrate these data sets to uncover biological insights. Here, we developed a domain-adversarial and variational approximation framework, DAVAE, to integrate multiple single-cell data across samples, technologies and modalities without any *post hoc* data processing. We fit normalized gene expression into a non-linear model, which transforms a latent variable of a lower-dimension into expression space with a non-linear function, a KL regularizier and a domain-adversarial regularizer. Results on five real data integration applications demonstrated the effectiveness and scalability of DAVAE in batch-effect removing, transfer learning, and cell type predictions for multiple single-cell data sets across samples, technologies and modalities. DAVAE was implemented in the toolkit package “scbean” in the pypi repository, and the source code can be also freely accessible at https://github.com/jhu99/scbean.

## Introduction

Single-cell sequencing technologies have emerged over the past decade as an extraordinarily sensitive technique that can quantatively measure gene expression levels [1], DNA methylation landscape [2], chromatin accessibility [3], in situ expression [4] at the level of single cell. Tremendous single-cell data sets are generated across different technologies, organisms, and modalities; and some large-scale comprehensive single-cell atlases [5, 6, 7] are now being built, which will cover almost every aspect of biology and complex diseases. Therefore, we are facing challenges of developing scalable and efficient methods for integrating single-cell data sets across samples, technologies and modalities; and gain biological insights into cellular heterogeneity, biological states/cell types, cell development and spatial patterns in complex tissues [8].

One major task in single-cell data integration is to remove various data noises, such as batch-effects, which hinder our ways of comparing two or more heterogeneous tissues. Both scmap [9] and scAlign [10] take the reference-based strategy to annotate query data sets, while these methods are unable to predict new cell types of query data sets. Batch-correction methods are proposed to overcome the deficiency, which attempt to remove batch effects from original datasets and return a batch-corrected expression matrix. Some previously existing batch-correction methods were specially designed for bulk RNA-seq, such as combat [11], RUVseq [12] and limma [13]. However, their applications to scRNA-seq is not pratical since their models assume that the cellular composition of each batch is identical. To avoid the assumption of equal composition, mnnCorrect [14] finds most similar cells by detecting mutually nearest neighbors (MNN) across batches, and obtains a batch correction vector by averaging many MNN pairs. Seurat [15] attempts to remove the batch effects on a set of metagene vectors using canonical correlation analysis (CCA) as an initial dimension reduction. Inspired by MNN, Seurat v3 [16] utilizes k-MNN for each cell within its paired dataset to identify matching pairs, termed as “anchors”, based on the CCA-based cell embeddings. However, its CCA-based dimensional reduction is potentially less efficient in capturing the structure of scATAC-seq data. That’s why it relies on latent semantic indexing to reduce the dimension for the scATAC-seq data. In analogy to mnnCorrect, Scanorama [17] employs a generalized mutual nearest-neighbors matching method to find similar cells among all datasets, instead of paired datasets, on lower dimensional embeddings. A probabilistic model scVI [18] based on a hierarchical Bayesian model with conditions was proposed to fit the count expression data into a zero-inflated negative binomial (ZINB) distribution conditioned on a batch annotation, as well as two additional unobserved random variables. LIGER [19] attempts to identify both shared cell types and dataset-specific features using integrative non-negative matrix factorization (iNMF).

Although several existing approaches (e.g. Seurat v3 and LIGER) provided efficient ways of integrating multiple scRNA-seq data sets, few of them can integrate single-cell data sets across modalities without *post hoc* data processing. Further, scalability and computation speed are essential problems when we integrated large-scale data sets. To address these limitations, we proposed a unified framework which can integrate multiple scRNA-seq into an atlas reference, transfer learning for scATAC-seq data, spatially resolved transcritomics without *post hoc* data processing.

## Results

### Methods overview

Here, we considered the problem of integrating multiple scRNA-seq data sets, and multiple single-cell data across modalities. To solve this problem, we proposed a unified framework, Domain-Adversarial and Variational Auto-Encoder (DAVAE), to fit the normalized gene expression (or chromatin accessibility) into a non-linear model, which transforms a latent variable *z* into the expression space with a non-linear function, a KL regularizier and a domain-adversarial regularizier. As shown in Fig. 1, DAVAE was implemented in the structure of neural networks, which consists of a variational approximation network [20], a generative Bayesian neural network and a domain-adversarial classifier [21]. The non-linear feature enables us to fit to complex models to capture the biological insights from the integration of multiple single-cell data sets; the deep neural network enables us to efficiently learn the regression model from large-scale data sets. In contrast to existing integration methods, it can transform multi-modalities into lower dimensional space without any *post hoc* data processing. The latent factors or variables in the shared-lower dimensional space can be used for clustering cell sub-populations, trajectory inference, and transfer learning across modalities, and the recovered expression data support downstream integrative analyses.

**Figure 1:**
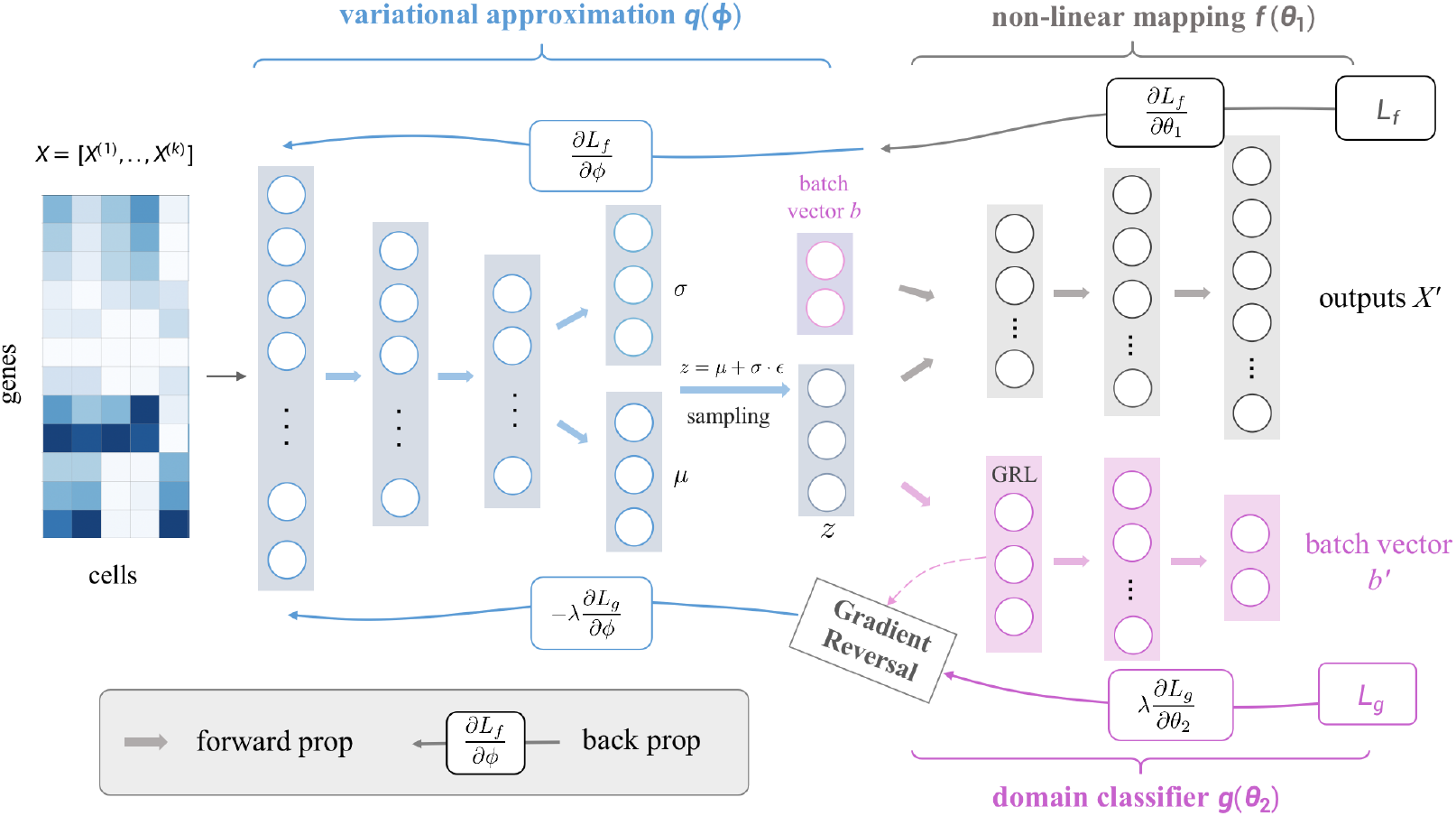
Overview of DAVAE. DAVAE is an adversarial and variational deep neural network framework for integrating multiple single-cell data sets, which includes a variational inference model (blue), a non-linear mapping (grey), a domain-adversarial classifier (pink). The gradient reversal layer (GRL) enables the adversarial mechanism, which takes the gradient from the subsequence level and changes its sign before passing it to the preceding layer.

### Integrating human dendritic cells from different samples

Our first application attempts to integrate scRNA-seq data on human blood dendritic cells [22] obtained from eight different samples with Smart-Seq2 protocol [23]. Following Tran et al. [24], we considered plates “P7”, “P8”, “P9”, “P10” as batch 1, and “P3”, “P4”, “P13”, “P14” as batch 2. Both of the two batches consist of 384 cells and a same set of 26,593 genes. Before integration, the two batches were coupled with nuisance factors such as batch-effects, and cells across the two batches are unable to be mixed together based on the raw single-cell data (Fig. 2a). The nuisance noise hinder our ways of downstream integration analysis such as clustering. Thus, our aim here is to remove batch-effects between the two batches, while preserving the true biological signal through the process of integration. DAVAE and three other integration algorithms, DESC, Scanorama and Seurat V3, were applied to integrate the two batches. To evaluate the integration quality, we first visualized the integration data using UMAP (Fig. 2b-e). The UMAP visualization show that DAVAE was the only algorithm that retained the heterogeneity of cell types and mixed the two batches in each cell type. Next, we used adjusted rand index (ARI) [25] to assess cell type purity and batch mixing. By following [26], ARI cell type and 1-ARI batch score were measured such that a higher value represents a better performance in clustering and batch mixing. The results show that DAVAE and DESC are the two best methods in terms of both ARI cell type and 1-ARI batch, which are significantly higher than that of Seurat V3 and Scanorama (Fig. 2f). Additionally, we performed kBET [27] test on each cell type over a range of neighborhood size that covers 5 to 25% of the sample size, with 100 replicates for each neighborhood size, to quantify batch effects between the two batches. An overall kBET acceptance rate was measured on the integrated data such that a higher value indicates a better performance of an integration method in removing batch effects. Compared to other integration methods, DAVAE obtained the highest overall kBET acceptance rate for each of the four cell types: CD1C DC, CD141 DC, plasmacytoid DC (pDC), and double negative cells (Fig. 2g). Overall, DAVAE facilitates the integration of scRNA-seq data collected across different samples.

**Figure 2:**
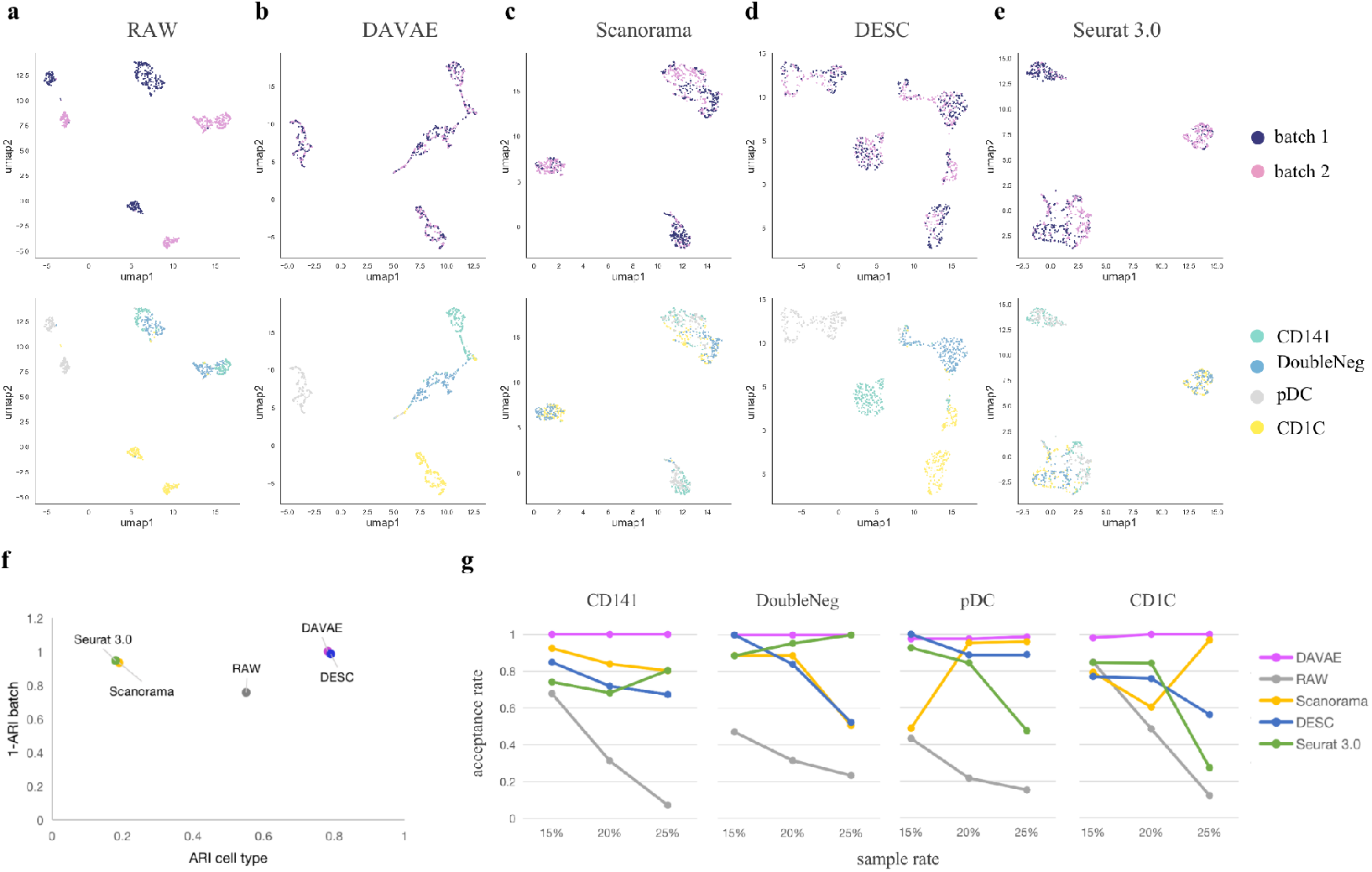
Integration of scRNA-seq data sets on human dendritic cells across samples. UMAP visualization on integrated data of five methods (RAW, DAVAE, Scanorama, DESC, Seurat 3.0) are organized into two rows (a-e). Each cell was represented by a dot and colored by batches (the first row), or cell types (the second row). Overall integration quality of compared algorithms are measured by two metrics: adjusted rand index (ARI) and kBET acceptance rate. Dot plots in (f) show clustering accuracy with ARI cell type and mixing quality with 1-ARI batch; Line plots in (g) show the average kBET acceptance rate of each integrated data across four cell types over a range of neighborhood size from 5 to 25% of the sample size.

### Integrating scRNA-seq data sets with different cellular compositions

Next, we examined our method in integrating three previously published scRNA-seq data sets [28]: one with 2,885 293T cells, one with 3,285 Jurkat cells, and one with a mixture of 1,605 293T cells and 1,783 Jurkat cells. It’s a challenge to remove batch effects without less- or over-correction for the first two data sets since their cell types are completely different. To verify the existence of batch-effects among the three data sets, we performed PCA on the raw scRNA-seq data before integration. UMAP visualization demonstrates that the Jurkat cells were separated into two distinct groups (Fig. 3a). To remove the batch-effects, we applied DAVAE, DESC, Scanorama, Seurat V3 to integrate these three data sets. Results show that DAVAE is the only method that can separate the three data sets into two different clusters and mix the same cell type well into the same cluster (Fig. 3b). DESC failed to separate these cells into two cell types; Scanorama and Seurat were unable to mix the same cell type well into a same cluster (Fig. 3c-e). Additionally, quantification in ARI cell type and 1-ARI batch suggest that DAVAE is superior to the three others in terms of clustering accuracy and batch mixing (Fig. 3f). DAVAE also achieves the highest kBET acceptance score for both of Jurkat cells and 293T cells with neighborhood settings from 5 to 25% of the sample size (Fig. 3g). Overall, DAVAE outperforms other existing methods in facilitating integration of data sets which may contain different cellular compositions.

**Figure 3:**
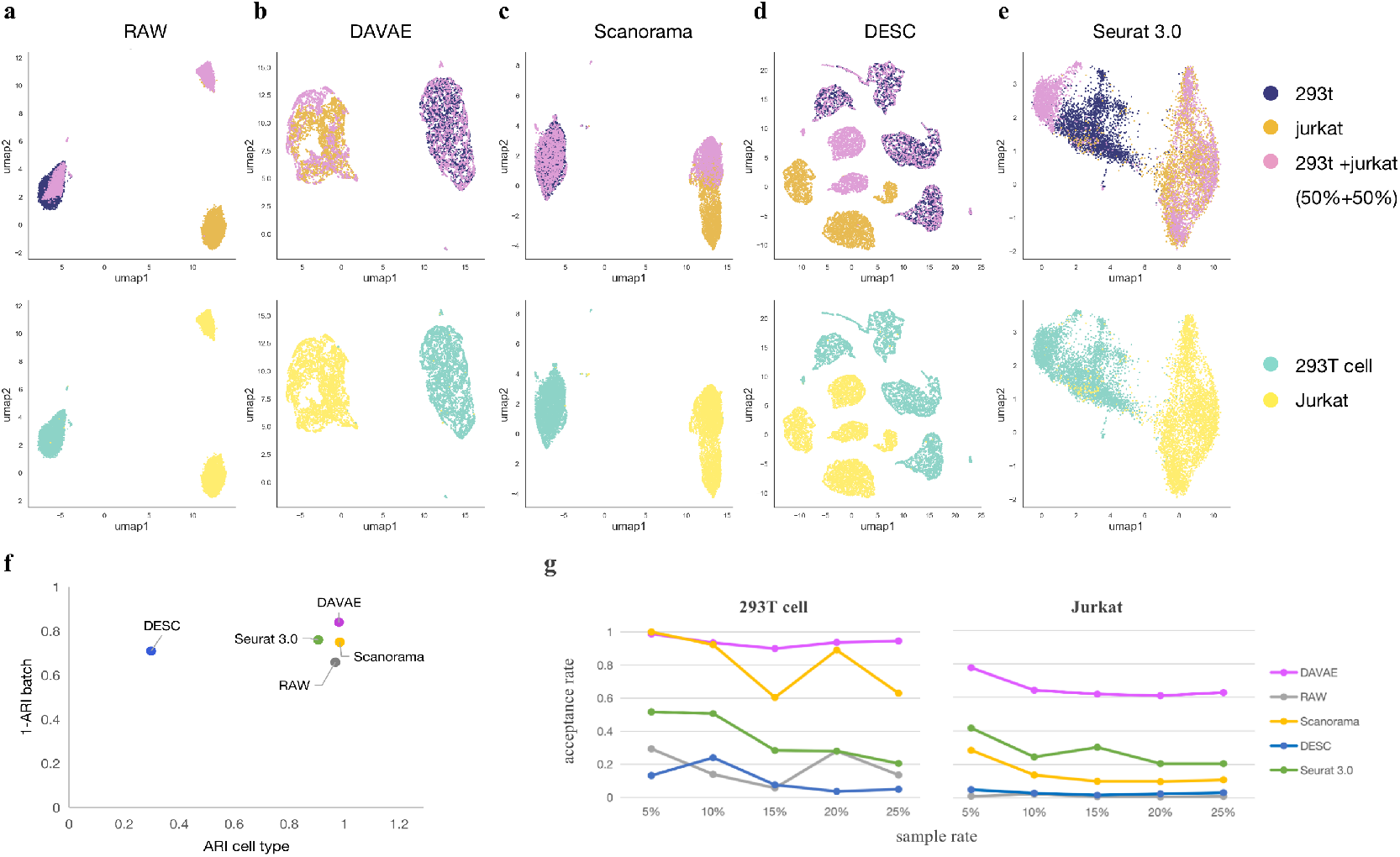
Integration of scRNA-seq data sets on 293T and Jurkat cells. UMAP visualization on integrated data of five methods (RAW, DAVAE, Scanorama, DESC, Seurat 3.0) are organized into two rows (a-e). Each cell was represented by a dot and colored by batches (the first row), or cell types (the second row). Overall integration quality of compared algorithms are measured by two metrics: adjusted rand index (ARI) and kBET acceptance rate. Dot plots in (f) show clustering accuracy with ARI cell type and mixing quality with 1-ARI batch; Line plots in (g) show the average kBET acceptance rate of each integrated data across two cell types over a range of neighborhood size from 5 to 25% of the sample size.

### Integrating scATAC-seq and scRNA-seq data on PBMCs

In our third application, we further extended DAVAE to integrate two different single-cell data types on human peripheral blood mononuclear cells (PBMCs): a scRNA-seq data of 33,538 genes measured on 11,769 cells and a scATAC-seq data of 78,700 peak measurements on 7,064 nuclei. Single cell ATAC-seq technology quantatively measures chromatin accessiblity in single cell resolution, which may provide new insights into cellular heterogeneity in tissue samples by linking cell type-specific regulatory variation and phenotypic variation [29]. Before integration with scRNA-seq data, we simply collapsed the peak matrix into an activity matrix by summing up all counts of peaks within a same region (a gene body and its 2k upstream). Then, we performed integration for the scRNA-seq data and the gene activity matrix, and successfully mixed the two data in the shared-lower dimensional space (Fig. 4a). Cell type labels of scATAC-seq cells were predicted by using a deep-learning-based classifier. The integrated scRNA-seq data (i.e. cell embedding) along with its cell type labels were treated as training data, or reference data, and each of the scATAC-seq cells was assigned to a predicted cell type. UMAP visualization shows that cells of different cell types were clearly separated into different groups in the latent feature space after integration performed by DAVAE (Fig. 4b). We performed kBET test on the cell embedding integrated by Seurat v3 and DAVAE over a range of neighbor size from 5 to 25% of the sample size. As shown in Fig. 4c, it suggests that cell embedding obtained by DAVAE is much better than that of Seurat v3. To verify the predicted cell type of scATAC-seq cells, we examined activity patterns of six well-known cell-type-specific marker genes: LYN, VCAN, CCL5, BANK1,SULF2, LDHB. Results in Fig. 4d suggests that our predicted cell types of scATAC-seq data were clearly separated into distinct groups. We further verified the predicted cell types by examining peak patterns within the gene body and the neighborhood of their corresponding marker genes (Fig. 4e). The expected chromatin accessibility patterns in the UMAP visualization and peaks of marker gene suggest that DAVAE can accurately transfer cell type labels from scRNA-seq data to scATAC-seq cells. On the other hand, we correctly failed to identify platelet cells in scATAC-seq data, because these cells are not nucleated [16]. Hence, we can draw a conclusion that DAVAE facilitates integration of scRNA-seq and scATAC-seq data and transfer learning for cell-type labels across modalities.

**Figure 4:**
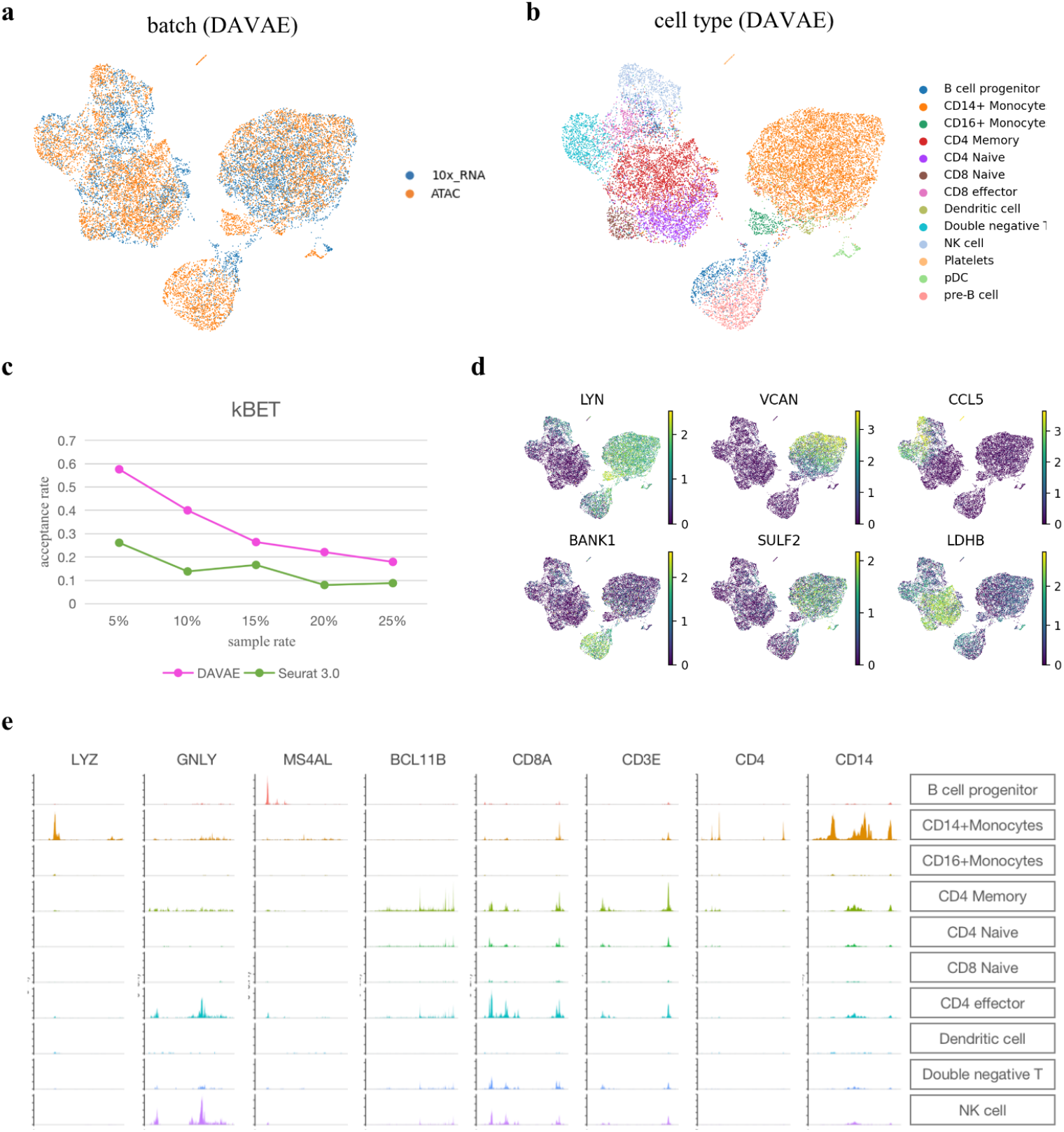
Integrating scATAC-seq data and scRNA-seq data on PBMCs. UMAP visualization shows DAVAE’s integrated results on scATAC-seq and scRNA-seq data. Each cell was represented by a dot and colored by batches (a) or cell types (b). Line plots in (c) show the average kBET acceptance rate of integrated data of Seurat v3 and DAVAE over a range of neighborhood size from 5 to 25% of the sample size. UMAP heatmaps in (d) show activity of six markers in scATAC-seq data, that include LYN, VCAN, CCL5, BANK1, SULF2, LDHB. Plots in (e) show peak activity patterns nearby the marker genes of ten cell types (B cell progenitor, CD14+ Monocyte, CD16+ Monocyte, CD4 Memory, CD4 Naive, CD8 Naive, CD4 effector, Dendritic cell, Double negative T, NK cell).

### Integrating spatial transcriptomics data and scRNA-seq data on mouse brain

We next extended our method to integrate two spatially resolved transcriptomics data sets, and used a technique of transfer learning to transfer labels from a reference of well-annotated scRNA-seq data to the spatial data. Specifically, we used two slices of mouse brain (anterior and posterior) sagittal data sets profiled with the 10X/Visium technology [30], and one scRNA-seq reference data profiled with smart-seq on Mouse Brain cortex with 22,272 cells on 36,577 genes [31]. Expression of 32,285 genes were measured for 2,695 spots in anterior and 3,355 spots in posterior. The Visium technology can measure gene expression profile, meanwhile, retain the background information of spatial coordinate in tissue samples. The integration of single-cell data of heterogeneous tissues across modalities aims to discover biological insights from spatially resolved transcriptomics with assistance of characterized scRNA-seq reference atlas. UMAP visualization shows that cells of the two slices were well mixed after integration (Fig. 5a). From the 26 clusters identified from the integrated data, we can clearly visualize the stratification of the cortical layer in both of the two tissues (Fig. 5b,c) in spatial coordinates. Additionally, the continuity between clusters in the two slices demonstrates that DAVAE can remove batch effects and preserve the tissue heterogeneity after the integration of spatial transcriptomics. To predict cell type labels for the Visium data, we further extended DAVAE to integrate the integrated Visium data and the scRNA-seq data across modalities, and performed a soft reference-based classification by assigning cell type-specific probabilities of each spot in the Visium data to each cell type. These assigned probabilities (weights) can reflect similarity between expression profile of spots in Visium data and expression profile of cell types in scRNA-seq data. From the heatmap results in spatial coordinate (Fig. 5d, Supplementary Fig. 1), we can see a clear successive layers of cortical neurons in both the anterior and posterior sagittal slices. Overall, DAVAE facilitates integrating not only two spatially resolved transcriptomics, but also scRNA-seq and spatially resolved transcriptomics across modalities.

**Figure 5:**
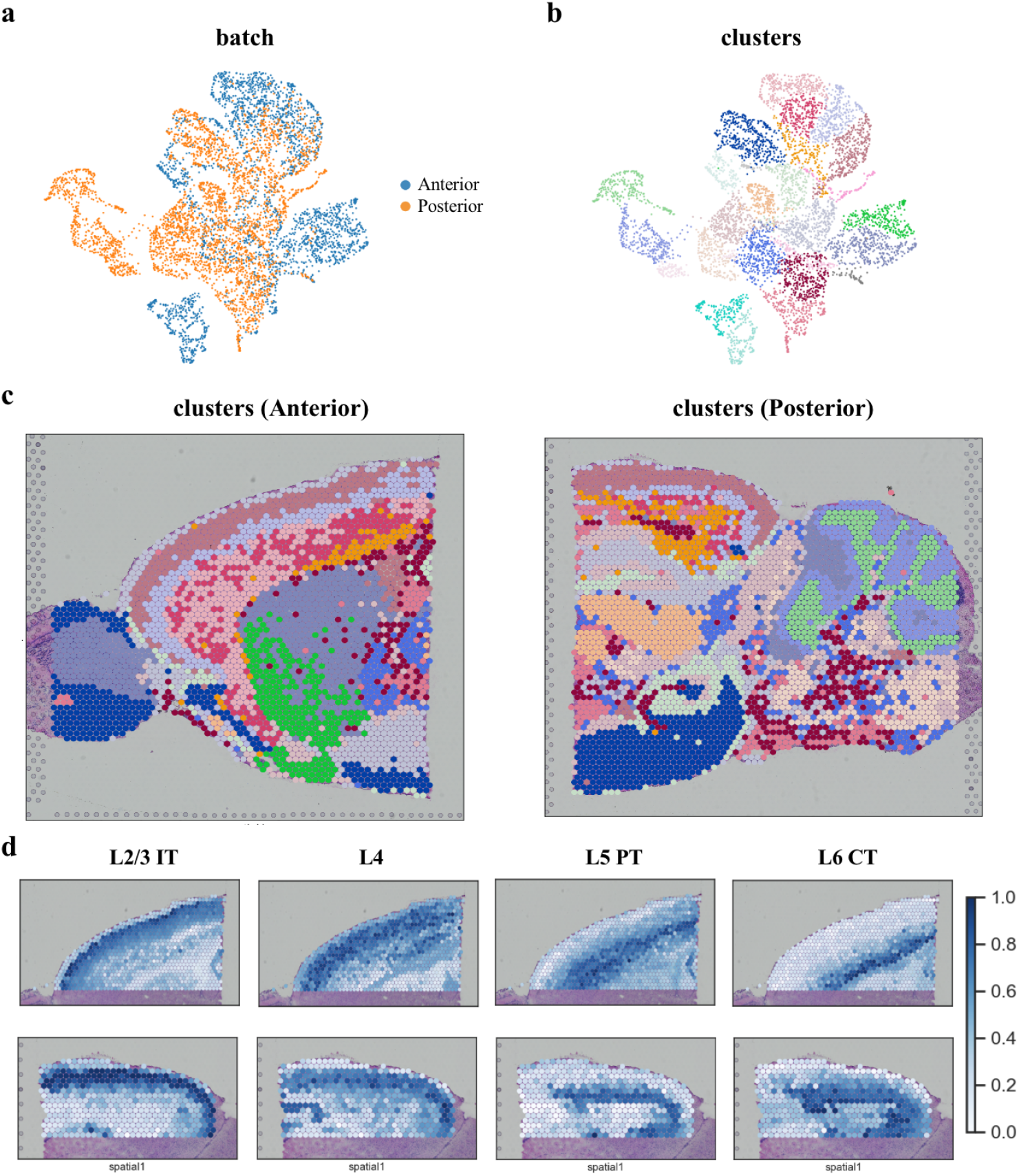
Integration of spatial transcriptomics data sets and scRNA-seq data set on sagittal anterior and posterior mouse brain. (a) UMAP visualization of mouse brain cells colored by batch label after DAVAE’s integration. (b) UMAP visualization of mouse brain cells colored by identified clusters after DAVAE’s integration. (c) Identified clusters in the spatial coordination. (d) After the integration of the spatial data with the scRNA-seq data on mouse cortex, cell-type-specific weight of each spot was learned from the reference scRNA-seq data. The heatmap plots show the probability of cell types L2/3IT, L4, L5PT, L6 CT in each spot. Predicted cell types were show in the supplementary figure.

### Integrating large-scale data sets

To test the scalability, DAVAE was applied to integrate two large-scale scRNA-seq data sets from human cell atlas (HCA): one with 384,000 cord blood cells; the other with 378,000 bone marrow cells [5]. Gene expression of each cell was measured on 18,969 genes with 10x genomics protocol. UMAP visualization shows that DAVAE can effectively mix the two data sets and separate each cell type into different clusters (Fig. 6a,b). Expression patterns of fourteen well-known marker genes support a similar conclusion that cell heterogeneity of the two batches was well preserved after integration performed by DAVAE (Fig. 6c-e). Specifically, heatmap of six marker genes (Fig. 6c) clearly show the six different groups of immune cells: B cell (CD79A); Dendritic (S100A8); Erythrocyte (HBB); NK cell (GNLY); and Myeloid (CST3); Memory T cells (CD3D); violin plot (Fig. 6d) and dot plot (Fig. 6e) show differential expression patterns of the fourteen marker genes across the ten clusters identified from the integrated data of DAVAE. The cell type annotation of each cluster was assigned based on the expression patterns of marker genes. To test the computational efficiency, we performed DAVAE, DESC and Scanorama on five sub-datasets uniformly down-sampled from the two HCA data sets with a machine of Intel(R) Xeon(R) Gold 6226R CPU @ 2.90GHz and 256G memory. The five data sets contain 100,000, 200,000, 400,000 and 600,000 cells, respectively. Results (Fig. 6f) show that deep learning-based algorithms (DAVAE and DESC) have significant advantages over Scanorama in terms of running time when applied to large-scale data sets. Overall, DAVAE is both efficient and scalable for integration of large-scale data sets.

**Figure 6:**
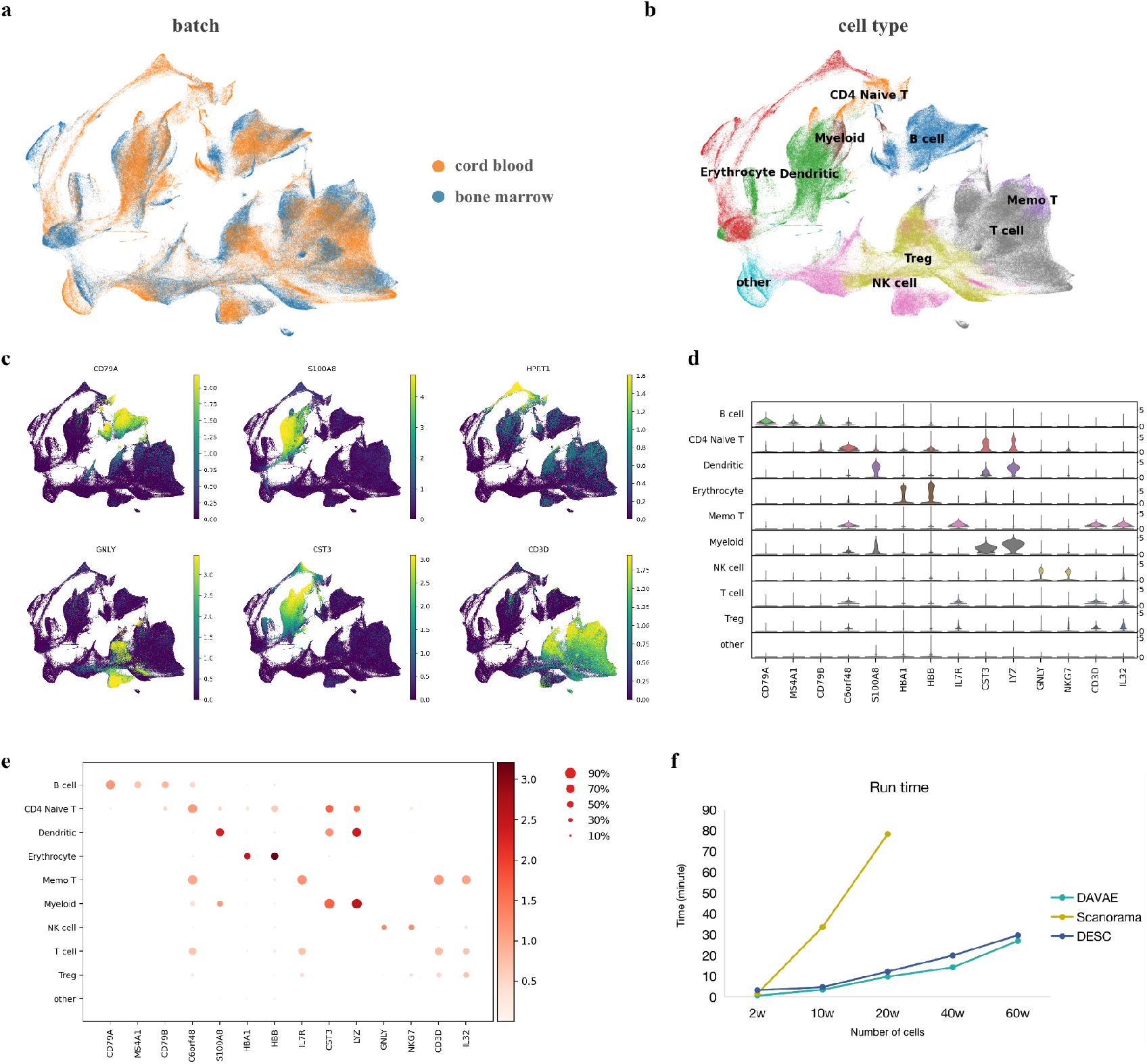
Integrating two large-scale data sets on human immune cells. UMAP visualization in (a) and (b) shows DAVAE’s integrated data on human cord blood and bone marrow. Each cell was represented by a dot and colored by batches (a) or cell types (b). Expression patterns of five marker genes (CD79A, S100A8, HPRT1, GNLY, CST3, CD3D) were shown in (c). Violin plots in (d) and dotplots in (e) show expression patterns of fourteen marker genes in ten cell types. Color in (d) represents the mean expression within each of the categories and the dot size in (e) indicates the fraction of cells in the categories expressing a gene. Line plots in (f) show runtime of DAVAE, DESC, Scanorama on five data sets with 100,000, 200,000, 400,000 and 600,000 cells, respectively.

## Discussion

In this paper, we proposed a novel non-linear model that consists of a non-linear function, a KL regulazier and a domain-adversarial regulazier to integrate multiple single-cell data sets across samples, technologies and modalities. To estimate the representation of cells in a commonly shared low-dimensional space, we constructed a variational and adversarial deep neural networks, called DAVAE, to unsupervisedly and jointly learn a variational approximation model, a generative model and a domain-adversarial classifier. DAVAE takes the normalized gene expression matrix as input and returns integrated data, which include cell embedding and recovered expression data that can be used for downstream integrative analyses, such as clustering, visualization, transfer learning across modalities. Comparing to existing methods, DAVAE can integrate multiple single-cell data sets across modalities without any *post hoc* data processing. To examine the effectiveness and scalability of DAVAE, we applied our methods and several widely-used existing approaches in integrating five real data sets. After a careful comparison, results demonstrate that DAVAE can effectively remove batch effects in multiple scRNA-seq data sets, while preserve the biological difference for various cell types. Further, our results demonstrate that DAVAE can integrate two consecutive spatial transcriptome sections to restore a complete tissue section; it can also integrate scATAC-seq data and spatial transcriptomics with a reference scRNA-seq data; and it can perform prediction of cell type labels using transfer learning across modalities. Lastly, by taking advantage of a mini-batch based stochastic gradient descent procedure, DAVAE is scalable and efficient for integration of large-scale scRNA-seq data sets, and can be accelerated by using GPUs. Our method is unsupervised and therefore does not require a prior knowledge about cell types. As single-cell genomics continues to evolve and sequencing experiments scale up, we believe that the efficient and scalable nature of DAVAE could make it a valuable tool for biomedical researchers to distinguish complex cellular heterogeneity.

## Methods and Materials

### The non-linear model

Let *X* = {*X*^(1)^, *X*^(2)^, · · · , *X*^(*k*)^} be *k* normalized gene expression matrices collected from *k* different scRNA-seq datasets with their corresponding batch-specific one-hot vectors {*b*^(1)^, *b*^(2)^, · · · , *b*^(*k*)^}. The *m*-th gene expression matrix *X*^(*m*)^ is a matrix of dimensionality *n_m_* by *p*, denoting normalized expression of *n_m_* cells on a common set of *p* genes across the *k* datasets. We extend our model to other single-cell data such as scATAC-seq, spatially resolved transcriptomics, in which *X*^(*m*)^ represents reads count of chromatin-accessible regions or counts of spatial spots. To integrate the *k* scRNA-seq datasets, it always involves finding a representation matrix *Z*^(*m*)^ of dimensionality *n_m_* by *p* for the expression matrix *X*^(*m*)^, where *d* ≪ *p* and *m* = 1, … , *k*. The set of lower-dimensional matrices *Z* = {*Z*^(1)^, *Z*^(2)^, · · · , *Z*^(*k*)^} are expected to reflect the true biological states of cells, and could be used for downstream analyses such as identification of cell subpopulations, trajectory inference, visualization. To estimate *Z*, we modeled the normalized gene expression into a non-linear model that transforms a latent variable *z* into expression space. Mathematically, we can write it as bellows:

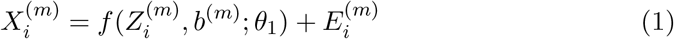

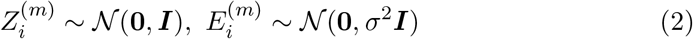

where 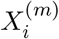 is a single-cell expression vector; *f* is a non-linear regression function that transforms 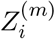 from a latent *d*-dimensional space into observable expression space with a one-hot vector *b*^(*m*)^ and a set of parameters *θ*_1_; and 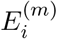 is a p-vector of residual errors following 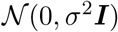. We assume that the latent factor 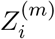 follows a standard multivariate normal distribution 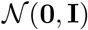. Under this assumption, the latent variables of different datasets all reside on the same *d*-dimensional space. The non-linear function *f* (·) was constructed by using a generative deep neural network structure, which takes 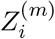, *b*^(*m*)^ as input, and outputs an outcome p-vector. In the deep neural networks, 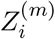 and *b*^(*m*)^ are concatenated into a (*d* + *k*)-dimensional layer. The resulting concatenated layer is connected to the output *p*-dimensional layer in the form of (*d* +*k*) → 32 → 64 → 128 → *p*. All intermediate layers are fully connected with each other through a batch normalization layer, relu activation function and a dropout layer (rate=0.1). Specifically, the batch normalization layer is designed to center and scale the inputs to a layer in a deep learning neural network, with the centering and scaling parameters treated as unknown and inferred through the inference algorithm. The dropout is a technique that can prevent over-fitting and provide a way of approximating combining exponentially many different neural network architectures efficiently. The final output layer is connected with *softplus* as activation function.

### Two regulariziers and the objective function

Our goal is to obtain estimates for 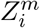 which can be used for downstream integrated analyses and visualization. However, the likelihood and the posterior probability of 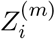 cannot be computed analytically because of non-linear function *f* (·). Thus, we propose to combine the above non-linear regression model, variational approximation approach [20] and a *domain-adversarial model* [21] within one process to jointly learn an inference model *q_ϕ_*(·), a domain-adversarial classifier 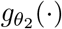 and the non-linear model 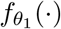. The main idea is to add two regularizers: KL regularizer and domain-adversarial regularizer, to the objective loss function.

#### KL regularizier

Mathematically, the marginal log-likelihood is composed of a sum over the marginal log-likelihood of individual cells, which can be written as:

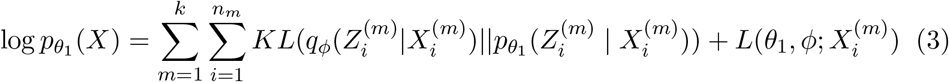

Here, the first term is KL divergence between the variational distribution and the posterior distribution; the second term is called evidence lower bound (ELBO). As KL divergence is non-negative, the MLE is equivalent to maximizing the ELBO. In other words, the ELBO hits the log probability of *X iff* the KL divergence is perfectly closed to 0. The ELBO can be written as:

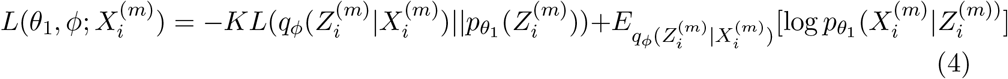

For simplicity, we assume that both the prior 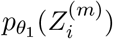 and the variational approximate posterior are Gaussian with a diagonal convariance:

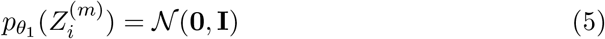

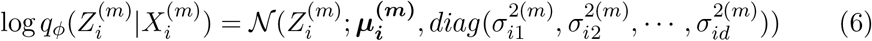

Then, the KL term in equation (4) can be computed analytically. To estimate the second term of equation (4), we used Monte Carlo estimator by sampling 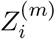 *N* times from 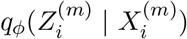. Since the residual errors follow 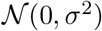, the second term can be rewritten as:

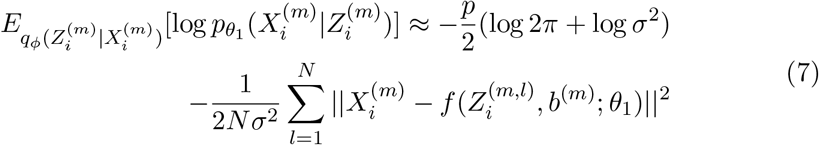

Let’s take the derivative with respect to *σ* and set it to zero. It yields the update rule:

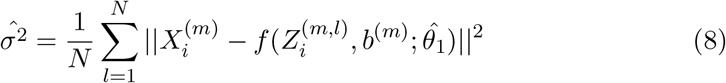

Now, the objective loss function is constrained with a KL regularizer, which can be written as belows:

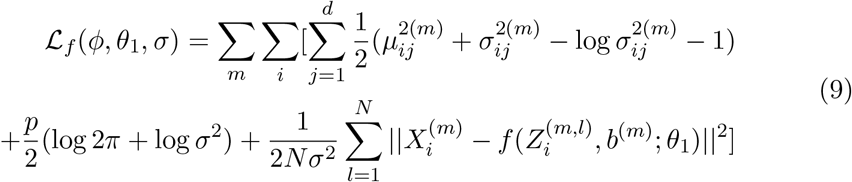

Here, 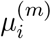 and 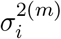 are estimated by learning the neural network. However, the Monte Carlo sampling process would result in indifferential bottleneck layer in the deep neural networks. To solve this problem, we used the reparameterization trick as follows:

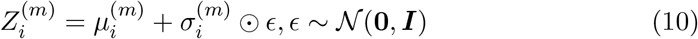

We rewrite the expectation and take samples of 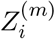 from the distribution *p*(*ϵ*) so that the sampling process is independent from the parameter 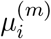 and 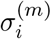. The symbol ⊙ represents element-wise multiplication of two vectors.

#### Domain-adversarial regularizier

Next, we embedded a domain-adversarial classifier 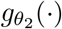 into the process of learning latent feature representation, so that the estimated latent factor 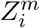 can represent biological state of cells across data sets and the original batch label cannot be classified by learning from its latent representation. The domain classifier is also constructed by using neural network structure, which takes 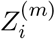 as input, outputs a probability distribution through one intermediate gradient reversal layers (GRL) and one dense layer. Specifically, the domain classifier are connected in the form of *d* → 16 → *k*. The *d* represents a *d*-dimensional GRL, which updates parameters with a reverse gradient using back-propagation gradient descent method. The GRL is fully connected with the following 16-dimensional dense layer through a relu activation function; and the final output *k* layer is supplied with softmax as the activation function and categorical cross entropy between the output probability and *b*^(*m*)^ as the loss function 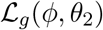.

#### The objective function

Now, we have constructed a non-linear model 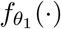, an inference model *q_ϕ_*(·) and a domain-adversarial learning model 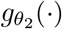 for expression vector 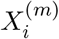 and a latent variable 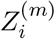. We can write the objective loss function as:

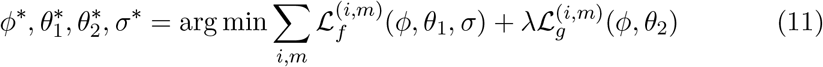

Then, our integration problem has been transformed into an optimization problem, which aims to search for (near-) optimal parameters by minimizing the objective function in equation (11). The hyper-parameter *λ* is used to tune the trade-off between these two quantities during the learning process. The gradient descent method updates each set of parameters in an iterative ways as below:

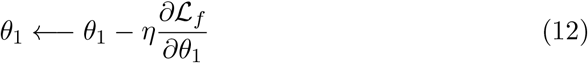

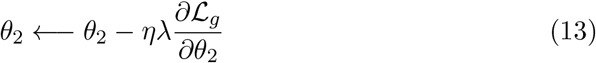

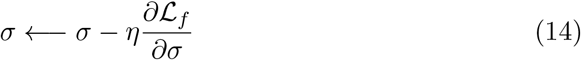

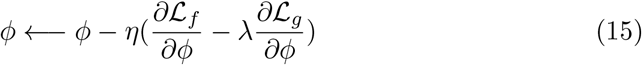

Here, *η* is the learning rate of gradient descent method. Note that the adversarial mechanism is achieved by minimizing 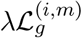 by updating parameters *θ*_2_ as 13, meanwhile maximizing 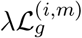 by updating parameters *ϕ* as 15 with GRL that takes the gradient from the subsequence level 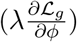 and changes its sign 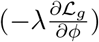 before passing it to the preceding layer.

### Data used in real data applications

#### Application 1: human dendritic cells

In the first application, human blood dendritic cell (DC) scRNA-seq data sets were obtained from GEO (GSE94820) [22], downloaded from https://www.ncbi.nlm.nih.gov/geo/query/acc.cgi?acc=GSE94820. We normalized the raw data with transcript per million (TPM) values. Cell type and batch information were extracted using the cell ID which is formatted as “Cell.Type”_“Plate.ID”. We considered plates “P7”, “P8”, “P9”, “P10” as batch 1, and “P3”, “P4”, “P13”, “P14” as batch 2. Both of the two batches consist of 384 cells and a same set of 26,593 genes.

#### Application2: 293T+ Jurkat cell

In the second application, we obtained three published data sets [28] from the 10X Genomics single-cell portal: one with 2,885 293T cells (https://support.10xgenomics.com/single-cell-gene-expression/datasets/1.1.0/293t); one with 3,285 Jurkat cells (https://support.10xgenomics.com/single-cell-gene-expression/datasets/1.1.0/jurkat), and one with a mixture of 1,605 293T cells and 1,783 Jurkat cells(https://support.10xgenomics.com/single-cell-gene-expression/datasets/1.1.0/jurkat:293t_50:50). All these data are measured on a common set of 32,738 genes. The cell type labels of the 50:50 mixed samples were obtained from 293t jurkat cluster.txt file in the cell labels subfolder of data downloaded from http://scanorama.csail.mit.edu/data.tar.gz

#### Application 3: human cell atlas

In our third application, we used data sets with 384,000 cord blood-derived cells in batch 1 and 378,000 bone marrow cells in batch 2 [5]. (cord blood: https://s3.amazonaws.com/preview-ica-expression-data/ica_cord_blood_h5.h5; bone marrow:https://s3.amazonaws.com/preview-ica-expression-data/ica_bone_marrow_h5.h5). Gene expression of each cell was measured on 18,969 genes with 10x genomics protocol.

#### Application 4: spatial transcriptomics data

In our fourth application, we used data sets consist of two slices (anterior and posterior) of the Mouse Brain Sagittal datasets profiled with the 10X/Visium technology. Anterior downloaded from: https://support.10xgenomics.com/spatial-gene-expression/datasets/1.1.0/V1_Mouse_Brain_Sagittal_Anterior. Posterior downloaded from: https://support.10xgenomics.com/spatial-gene-expression/datasets/1.1.0/V1_Mouse_Brain_Sagittal_Posterior.

#### Application 5: scATAC-seq data

In our fifth application, we used datasets consist of two different single-cell data types on human peripheral blood mononuclear cells (PBMCs): a scRNA-seq data of 33538 genes measured on 11769 cells (https://support.10xgenomics.com/single-cell-gene-expression/datasets/3.0.0/pbmc_10k_v3) and a single cell ATAC-seq data of 78700 peak measurements on 7064 nuclei, both of which were profiled with 10X genomics technologies (https://cf.10xgenomics.com/samples/cell-atac/1.0.1/atac_v1_pbmc_10k/atac_v1_pbmc_10k_filtered_peak_bc_matrix.h5). We obtained 13 cell types in the scRNA-seq data using the standard work-flow in Seurat (https://www.dropbox.com/s/3f3p5nxrn5b3y4y/pbmc_10k_v3.rds?dl=1).

## Software availability

The DAVAE algorithm was implemented in our python package *scbean* (≥0.4.0), which was archived at https://github.com/jhu99/scbean. A tutorial that descibes how to use DAVAE for several real data applications was published at https://scbean.readthedocs.io/en/latest/?badge=latest. All source code and data sets used for reproducing our results have been deposited at https://github.com/jhu99/davae_paper.

## Acknowledgements

Publication costs were funded by the National Natural Science Foundation of China (Grant No. 62072374); This project has been funded by the National Natural Science Foundation of China (Grant No. 61332014 and 61772426); the China Postdoctoral Science Foundation (Grant No. 2017M613203).

## Author’s contributions

J.H. conceived the idea and provided funding support. J.H. and Y.Z. jointly designed the experiments and developed the method, implemented the software, wrote the manuscript. Y.Z. performed applications on real data. X.S. contributed many discussion for improving the manuscript.

## References

[1] Tang F, Barbacioru C, Wang Y, Nordman E, Lee C, Xu N, et al. mRNA-Seq whole-transcriptome analysis of a single cell. Nature methods. 2009;6(5):377–382.

[2] Luo C, Rivkin A, Zhou J, Sandoval JP, Kurihara L, Lucero J, et al. Robust single-cell DNA methylome profiling with snmC-seq2. Nature Communications. 2018;9(1):3824.

[3] Cao J, Cusanovich DA, Ramani V, Aghamirzaie D, Pliner HA, Hill AJ, et al. Joint profiling of chromatin accessibility and gene expression in thousands of single cells. Science. 2018;361(6409):1380–1385. Available from: https://science.sciencemag.org/content/361/6409/1380.

[4] Moffitt JR, Hao J, Wang G, Chen KH, Babcock HP, Zhuang X. High-throughput single-cell gene-expression profiling with multiplexed error-robust fluorescence in situ hybridization. Proceedings of the National Academy of Sciences. 2016;113(39):11046–11051. Available from: https://www.pnas.org/content/113/39/11046.

[5] Regev A, Teichmann SA, Lander ES, Amit I, Benoist C, Birney E, et al. Science forum: the human cell atlas. Elife. 2017;6:e27041.

[6] Han X, Wang R, Zhou Y, Fei L, Sun H, Lai S, et al. Mapping the mouse cell atlas by microwell-seq. Cell. 2018;172(5):1091–1107.

[7] Cusanovich DA, Hill AJ, Aghamirzaie D, Daza RM, Pliner HA, Berletch JB, et al. A single-cell atlas of in vivo mammalian chromatin accessibility. Cell. 2018;174(5):1309–1324.

[8] Shapiro E, Biezuner T, Linnarsson S. Single-cell sequencing-based technologies will revolutionize whole-organism science. Nature Reviews Genetics. 2013;14(9):618–630.

[9] Kiselev VY, Yiu A, Hemberg M. scmap: projection of single-cell RNA-seq data across data sets. Nature methods. 2018;15(5):359–362.

[10] Johansen N, Quon G. scAlign: a tool for alignment, integration, and rare cell identification from scRNA-seq data. Genome biology. 2019;20(1):1–21.

[11] Johnson WE, Li C, Rabinovic A. Adjusting batch effects in microarray expression data using empirical Bayes methods. Biostatistics. 2007;8(1):118–127.

[12] Risso D, Ngai J, Speed TP, Dudoit S. Normalization of RNA-seq data using factor analysis of control genes or samples. Nature biotechnology. 2014;32(9):896–902.

[13] Ritchie ME, Phipson B, Wu D, Hu Y, Law CW, Shi W, et al. limma powers differential expression analyses for RNA-sequencing and microarray studies. Nucleic acids research. 2015;43(7):e47–e47.

[14] Haghverdi L, Lun ATL, Morgan MD, Marioni JC. Batch effects in single-cell RNA-sequencing data are corrected by matching mutual nearest neighbors. Nature Biotechnology. 2018;36(5):421–427.

[15] Butler A, Hoffman P, Smibert P, Papalexi E, Satija R. Integrating single-cell transcriptomic data across different conditions, technologies, and species. Nature biotechnology. 2018;36(5):411–420.

[16] Stuart T, Butler A, Hoffman P, Hafemeister C, Papalexi E, Mauck III WM, et al. Comprehensive integration of single-cell data. Cell. 2019;177(7):1888–1902.

[17] Hie B, Bryson B, Berger B. Efficient integration of heterogeneous single-cell transcriptomes using Scanorama. Nature Communications. 2019;37(6):685–691.

[18] Lopez R, Regier J, Cole MB, Jordan MI, Yosef N. Deep generative modeling for single-cell transcriptomics. Nature methods. 2018;15(12):1053–1058.

[19] Welch JD, Kozareva V, Ferreira A, Vanderburg C, Martin C, Macosko EZ. Single-cell multi-omic integration compares and contrasts features of brain cell identity. Cell. 2019;177(7):1873–1887.

[20] Kingma DP, Welling M. Auto-encoding variational bayes. arXiv preprint arXiv:13126114. 2013;.

[21] Ganin Y, Ustinova E, Ajakan H, Germain P, Larochelle H, Laviolette F, et al. Domain-Adversarial Training of Neural Networks. 2015;.

[22] Villani AC, Satija R, Reynolds G, Sarkizova S, Shekhar K, Fletcher J, et al. Single-cell RNA-seq reveals new types of human blood dendritic cells, monocytes, and progenitors. Science. 2017;356(6335).

[23] Picelli S, Björklund ÅK, Faridani OR, Sagasser S, Winberg G, Sandberg R. Smart-seq2 for sensitive full-length transcriptome profiling in single cells. Nature methods. 2013;10(11):1096–1098.

[24] Tran HTN, Ang KS, Chevrier M, Zhang X, Lee NYS, Goh M, et al. A benchmark of batch-effect correction methods for single-cell RNA sequencing data. Genome Biology. 2020;21(1):12.

[25] Hubert L, Arabie P. Comparing partitions. Journal of Classification. 1985;2(1):193–218.

[26] Tran HTN, Ang KS, Chevrier M, Zhang X, Lee NYS, Goh M, et al. A benchmark of batch-effect correction methods for single-cell RNA sequencing data. Genome Biol. 2020 01;21(1):12.

[27] Büttner M, Miao Z, Wolf FA, Teichmann SA, Theis FJ. A test metric for assessing single-cell RNA-seq batch correction. Nature Methods. 2019;16(1):43–49.

[28] Zheng GXY, Terry JM, Belgrader P, Ryvkin P, Bent ZW, Wilson R, et al. Massively parallel digital transcriptional profiling of single cells. Nature Communications. 2017;8(1):14049.

[29] Buenrostro JD, Wu B, Litzenburger UM, Ruff D, Gonzales ML, Snyder MP, et al. Single-cell chromatin accessibility reveals principles of regulatory variation. Nature. 2015;523(7561):486–790.

[30] Ståhl PL, Salmén F, Vickovic S, Lundmark A, Navarro JF, Magnusson J, et al. Visualization and analysis of gene expression in tissue sections by spatial transcriptomics. Science. 2016;353(6294):78–82. Available from: https://science.sciencemag.org/content/353/6294/78.

[31] Tasic B, Yao Z, Graybuck LT, Smith KA, Nguyen TN, Bertagnolli. Shared and distinct transcriptomic cell types across neocortical areas. Nature. 2018;563(7729):72–78.

