## supplementary figures and text for "Efficient and scalable integration of single-cell data using domain-adversarial and variational approximation"

#### 1 Supplementary Figures

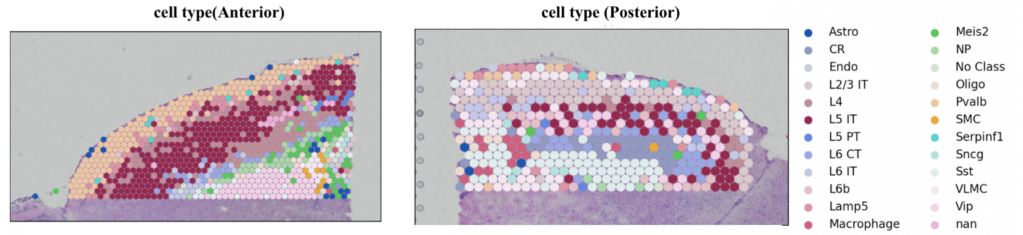


Figure 1: Transferring cell type labels to each spot of spatially resolved transcriptomics based on cell type knowledge of the reference scRNA-seq data. To do so, we first integrating spatial data and scRNA-seq data on mouse cortex. Then we performed reference-based classification to assign each spot a cell-type-specific probability. Lastly, the cell type of the highest probability was assigned to a spot.

### 2 Supplementary Text

#### Label transfer

We use a deep neural network classifier to transfer labels from scRNA-seq data sets of known cell types to other data sets. The structure of our classifier is a four-layer neural network, in which the output layer uses the Softmax activation function, and the remaining layers use the Relu activation function. The classifier takes the embedded features as input and the output is a vector $\hat{y} = (y_1, \ldots, y_i, \ldots, y_k)$, where each value $y_i$ represents the probability of a cell belonging to category $i$. The loss function of this classifier is the categorical cross entropy between $\hat{y} = (y_1, y_2, \ldots, y_k)$ and one-hot encoding $y$ of cell type label.

#### Data normalization and gene selection

At first, we perform CPM-normalization, where the UMI count of each gene in each cell is divided by the total number of UMI in the cell, multiplied by 1000,000. Then CPM-normalized data was converted to a natural logarithmic scale. This step was implemented by normalize_total and log1p function from the Scanpy package `https://github.com/theislab/scanpy`. Then we identify a subset of genes that exhibit high variability in the cell, thereby representing heterogeneous features for downstream analysis. High variable genes were selected using the highly_variable_genes function from the Scanpy package.

#### Evaluation metric for clustering

For published data sets in which the reference cell-type labels are known, we use ARI (Adjusted Rand Index) to compare the performance of different clustering algorithms. ARI is a measure of similarity between the clustering labels and the reference cluster labels. To assess cell type purity using ARI, we use k-means algorithm to cluster the integrated data and calculate the ARI between k-means cluster and reference cell-type labels through the sklearn package(`https://scikit-learn.org/stable/`). Given the contingency table 2, each value $n_{ij}$ in the table represents the number of a document in cluster $Y$ and class $X$ at the same time, and the ARI value can be calculated as follow:

$$ARI = \frac{\sum_{ij}\binom{n_{ij}}{2} - [\sum_i \binom{a_i}{2} \sum_j \binom{b_j}{2}]/\binom{n}{2}}{\frac{1}{2}[\sum_i \binom{a_i}{2} + \sum_j \binom{b_j}{2}] - [\sum_i \binom{a_i}{2} \sum_j \binom{b_j}{2}]/\binom{n}{2}} \tag{1}$$

| | $Y_1$ | $Y_2$ | ... | $Y_s$ | *Sums* |
|---|---|---|---|---|---|
| $X_1$ | $n_{11}$ | $n_{12}$ | ... | $n_{1s}$ | $a_1$ |
| $X_2$ | $n_{21}$ | $n_{22}$ | ... | $n_{2s}$ | $a_2$ |
| ... |  |  |  |  | ... |
| $X_r$ | $n_{r1}$ | $n_{r2}$ | ... | $n_{rs}$ | $a_r$ |
| *Sums* | $b_1$ | $b_2$ | ... | $b_s$ | |

$ARI \in [-1, 1]$, the larger the value means that the clustering result is more consistent with the real situation.

#### Evaluation metric for batch effect

*kBET* (k-nearest-neighbor batch estimation): We use kBET to quantify batch effects in scRNA-seq data. kBET uses a $\chi^2$-based test for random neighborhoods of fixed size to determine whether they are well mixed and its null hypothesis is"all batches are well-mixed". First, the algorithm creates k-nearest neighbor matrix and chooses 10% (sample rate, can be adjusted) of the samples to check the batch label distribution in its neighborhood. If the local batch label distribution is sufficiently similar to the global batch label distribution, the $\chi^2$-test does not reject the null hypothesis. The neighborhood size k is fixed for all tests. The test returns a binary result for each of the tested samples and the final result of kBET is the average test rejection rate. The lower the test result, the less bias is introduced by the batch effect which means superior mixing of data. kBET is very sensitive to any kind of bias. kBET was calculated by R package kbet (`https://github.com/theislab/kBET`).

*ARI-batch*: For batch mixing assessment, we also calculate the ARI between respective batch labels and the k-means clustering labels (ARI-batch), and a low ARI-batch score denotes superior mixing.
